# Intracellular defensive symbiont is culturable and capable of transovarial, vertical transmission

**DOI:** 10.1101/2023.12.05.570145

**Authors:** Gerald P. Maeda, Mary Katherine Kelly, Aadhunik Sundar, Nancy A. Moran

## Abstract

Insects frequently form heritable associations with beneficial bacteria that are vertically transmitted from parent to offspring. Long term vertical transmission has repeatedly resulted in genome reduction and gene loss rendering many such bacteria incapable of independent culture. Among aphids, heritable endosymbionts often provide a wide range of context-specific benefits to their hosts. Although these associations have large impacts on host phenotypes, experimental approaches are often limited by an inability to independently cultivate these microbes. Here, we report the axenic culture of *Candidatus* Fukatsuia symbiotica strain WIR, a heritable bacterial endosymbiont of the pea aphid, *Acyrthosiphon pisum*. Whole genome sequencing revealed similar genomic features and high sequence similarity to previously described strains, suggesting the cultivation techniques used here may be applicable to *Ca*. F. symbiotica strains from distantly related aphids. Microinjection of the isolated strain into uninfected aphids revealed that it can reinfect developing embryos, and is maintained in subsequent generations via transovarial maternal transmission. Artificially infected aphids exhibit similar phenotypic and life history traits compared to native infections, including protective effects against an entomopathogenic *Fusarium* species. Overall, our results show that *Ca*. F. symbiotica may be a useful tool for experimentally probing the molecular mechanisms underlying heritable symbioses and antifungal defense in the pea aphid system.

**IMPORTANCE:** Diverse eukaryotic organisms form stable, symbiotic relationships with bacteria that provide benefits to their hosts. While these associations are often biologically important, they can be difficult to probe experimentally, because intimately host-associated bacteria are difficult to access within host tissues, and most cannot be cultured. This is especially true of the intracellular, maternally inherited bacteria associated with many insects, including aphids. Here, we demonstrate that a pea aphid-associated strain of the heritable endosymbiont, *Candidatus* Fukatsuia symbiotica, can be grown outside of its host using standard microbiology techniques, and can readily re-establish infection that is maintained across host generations. These artificial infections recapitulate the effects of native infections making this host-symbiont pair a useful experimental system. Using this system, we demonstrate that *Ca*. F. symbiotica infection reduces host fitness under benign conditions, but protects against a previously unreported fungal pathogen.

## INTRODUCTION

Diverse insect lineages have independently evolved mechanisms that ensure stable, vertical transmission of beneficial microbes to offspring (1–4). Many of these heritable endosymbionts supplement nutrients lacking in the host diet and are essential for normal host growth and development (5–8). Others drastically alter host phenotypes, with some known examples influencing host thermal tolerance, pathogen or parasitoid resistance, body color, and dietary breadth (9–13). Despite the ubiquity and importance of these symbiotic relationships, intimate host association often presents practical challenges for experimental studies. Restriction to host tissues can present challenges in generating sequencing coverage for quality genome assemblies of endosymbionts present at low abundances. Most tools for forward or reverse genetics in microbes are designed for pure cultures, and cannot be performed on bacteria within host tissues. Most studied maternally transmitted endosymbionts can not be independently cultured using conventional microbiological techniques (14).

Aphids are globally distributed pest insects of the order Hemiptera that have been established as a useful model for understanding heritable symbioses. Nearly all aphids harbor an obligate endosymbiont, *Buchnera aphidicola,* that supplies essential amino acids lacking in the host diet (5, 6, 15). The association between *Buchnera* and aphids is ancient and over the course of an estimated 200 MY, *Buchnera* has experienced extreme gene loss and genome reduction (16). Several non-essential or facultative endosymbionts have formed more recent associations and can be found at intermediate frequencies in natural populations (17, 18). Though more recent associations, the process of maternal transmission for *Buchnera* and other facultative endosymbionts is similar, occurring early in embryonic development, generally results in stable transmission to all offspring of infected individuals (19, 20). Very early during embryonic development, *Buchnera* and any co-infecting facultative endosymbionts are endocytosed at the posterior end of the blastula, entering a central, syncytial cell, before packing into bacteriocytes. These facultative symbioses are more recent associations, and potentially more amenable for experimental manipulation. Culture-assisted methods would greatly facilitate our understanding of the molecular mechanisms underlying symbiont transmission in this system. However, to date, no heritable aphid endosymbionts have been successfully reintroduced and stably maintained after axenic culture.

One particularly promising candidate for axenic cultivation is the pea aphid endosymbiont, *Candidatus* Fukatsuia symbiotica (previously X-type or PAXS). So far, one pea aphid-associated strain, *Ca*. Fukatsuia symbiotica strain 5D, has been co-cultured with insect cells (21). This strain has relatively intact metabolic capabilities compared to other vertically transmitted aphid endosymbionts, has experienced only moderate gene loss, and has intermediate genome-wide GC content, suggesting a comparatively recent transition to a vertically transmitted lifestyle (21). *Ca.* F. symbiotica strains have variable effects on host phenotypes, with reported benefits ranging from heightened parasitoid resistance, protection against fungal pathogens, and increased tolerance to extreme heat (22–24). However, high costs to host fitness have also been observed in infected aphids, with some strains providing minimal relief to biotic or abiotic stressors (25). Given the lack of easily culturable, vertically transmitted bacterial symbiont, we attempted to axenically cultivate *Ca.* F. symbiotica from a naturally infected pea aphid, in an effort to establish a tractable system to study its diverse effects on the host.

## RESULTS

### *Candidatus* F. symbiotica strain WIR is capable of axenic culture

We attempted to grow *Ca*. *F. symbiotica* axenically from a naturally infected pea aphid using standard microbiology media. Growth is observed on heart infusion agar supplemented with 5% defibrinated sheep’s blood, with clear to cloudy colonies forming after 2 weeks of incubation at room temperature under ambient atmosphere. No growth was observable at higher temperatures (25 C, 30 C, or 35 C). Light microscopy revealed long (∼5 μm), and rod-shaped morphology (Fig. 1A). Scanning electron microscopy also confirmed long, rod-shaped morphology (Fig. 1B, C).

**FIG 1.**
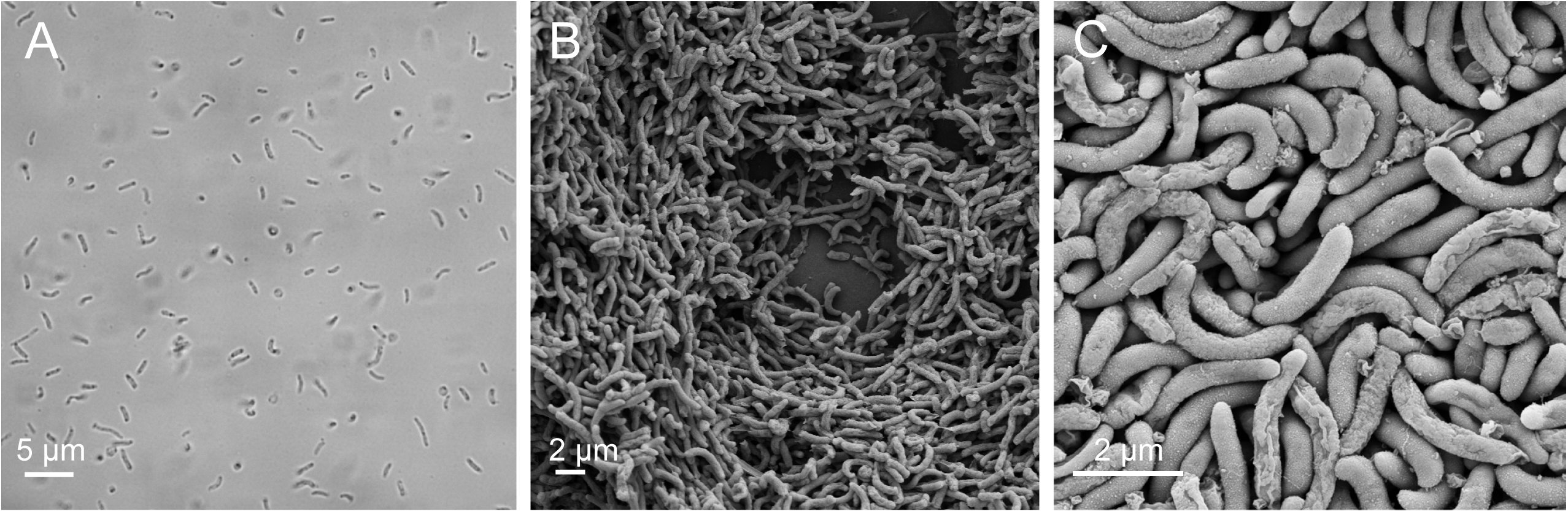
Morphology of *Ca.* F. symbiotica WIR cultivated outside of its aphid host. (A). Light microscopy of *Ca.* F. symbiotica WIR resuspended after growth on HIA with 5% sheep’s blood. Scale bar indicates 5 μm (B, C) Scanning electron microscopy (SEM) performed on plate-grown *Ca.* F. symbiotica WIR. Scale bars indicate 2 μm.

### Genomic features and phylogenetic placement of *Ca*. F. symbiotica WIR

*Ca*. F. symbiotica str. WIR was subjected to whole genome sequencing, using a combination of Illumina short reads and Oxford Nanopore long reads, for a total coverage depth of 463x. The complete genome assembly of *Ca*. F. symbiotica WIR has a total size (3.065 Mbp) and GC content (43.77%) very close to that of strain 5D (Table S1).

We generated a maximum likelihood phylogeny based on 340 shared single-copy orthologs, for phylogenetic placement of strain WIR. *Ca*. Fukatsuia strains form a well supported monophyletic clade within the family Yersiniaceae (Fig. 2A). Strains associated with pea aphid, *Cinara confinis*, and *Drepanosiphum platanoidis*, form a tight clade with the recently described strains associated with the aphid genus *Anoecia* forming a distinct sister clade consistent with recent work (26). Pairwise comparisons show that average nucleotide identity is also high (>99.8%) among the pea aphid, *Cinara*, and *Drepanosiphum* associated strains (Fig. 2B) with a steep decline in comparisons with *Anoecia*-associated strains.

**FIG 2.**
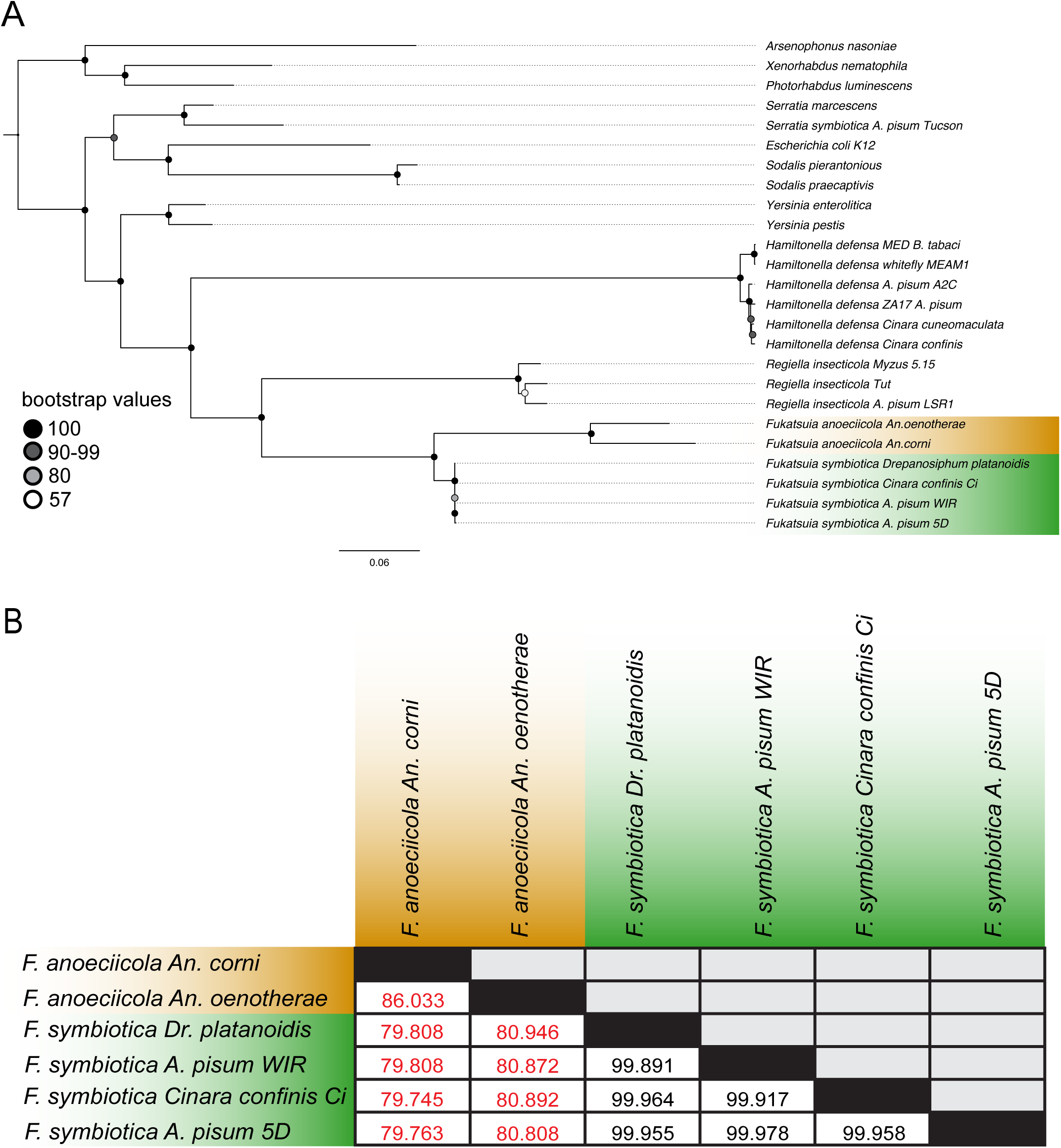
Phylogenetic placement and sequence similarity of *Ca*. F. symbiotica WIR relative to other strains of interest. (A) Maximum likelihood phylogeny of *Ca.* Fukatsuia strains and other Gram-negative bacteria, based on the concatenated amino acid alignments of 340 single copy orthologs. Scale bar indications number of substitutions per site. Genome accessions and related information are included in Table S2. (B) Pairwise comparisons of average nucleotide identity of *Ca*. Fukatsuia strains.

### Cultured *Ca*. F. symbiotica WIR is vertically transmitted to offspring following injection

Given its overall high genomic similarity to vertically transmitted strains and its stable maintenance in naturally infected aphids in the laboratory, we tested whether our *Ca*. F. symbiotica strain WIR is capable of stably recolonizing uninfected aphids. Bacterial colonization of aphid embryos is a selective process, only occurring during a specific stage early in embryonic development (19, 20). When transferring endosymbionts through hemolymph injection, only aphid embryos at the receptive developmental stage are colonized, resulting in a delay between time of injection and production of infected offspring. Among facultative endosymbionts that persist and proliferate within the insect hemocoel, this results in an increase in infection frequency with time from injection (27–29).

To determine if our cultivated *Ca.* F. symbiotica is capable of colonizing developing embryos after axenic culture, uninfected aphids were injected with a suspension of bacterial cells. Offspring produced as early as 8 days after injection tested positive for *Ca*. F. symbiotica infection (Fig. 3A). A general upward trend was observed, with 69% of offspring born 13 days after injection, testing positive. To determine if these experimentally established infections are stably maintained, a second set of injections were performed, using the same methods. Offspring born 10-12 days post-injection were screened for infection. Of the 24 aphids tested for *F. symbiotica*, 15 tested positive for infection (62.5%). These aphids were allowed to mature to adulthood and reproduce before sampling, allowing us to screen subsequent generations (Fig 3B). By the third generation after injection, 100% infection frequency was observed (Fig 3B). Six sublines of infected aphids were maintained by transferring three nymphs to new plants, biweekly. *Ca*. Fukatsuia symbiotica infection status was reexamined approximately 5 months later in these lines corresponding to approximately 10 generations after injection, and was detected in 100% (24/24) of the aphids screened. Thus, once established, *Ca*. Fukatsuia symbiotica is inherited with very high fidelity.

**FIG 3.**
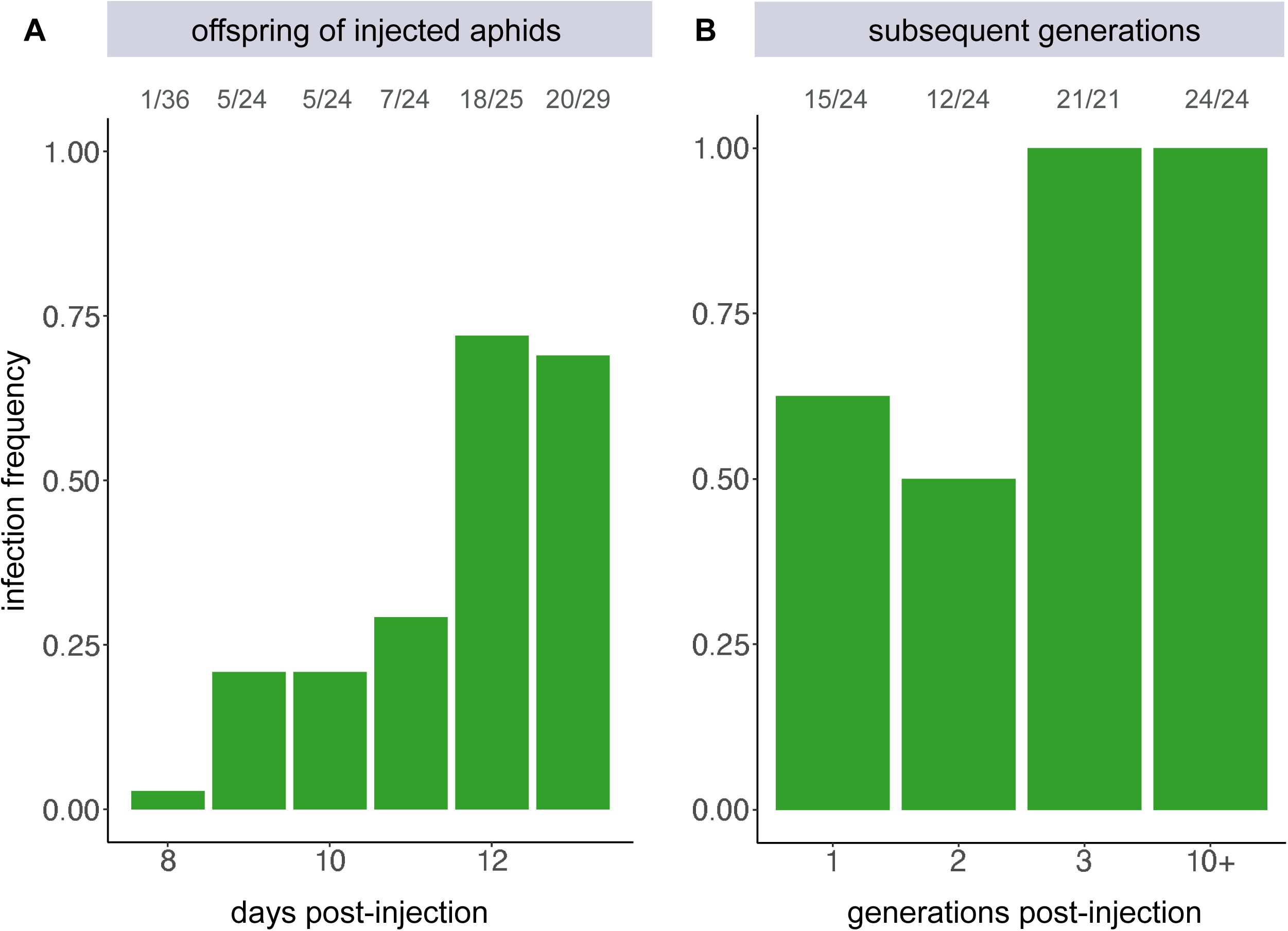
Colonization pattern of *Ca*. F. symbiotica WIR following injection into uninfected aphids. (A) Bar chart indicating the portion of individuals testing positive for *Ca.* F. symbiotica infection, in the generation immediately following injection, sampled daily. (B) Bar chart showing the portion of infected individuals in subsequent generations. The number of individuals testing positive over number sampled is shown above each bar.

### *Ca*. F. symbiotica WIR is maintained via transovarial transmission

To determine if the artificially generated *Ca.* F. symbiotica infections are maintained via the route of transovarial transmission known for other aphid symbionts (20, 28) fluorescent *in situ* hybridization (FISH) microscopy was performed. Three generations after injection, developing embryos were dissected out of adult aphids. These embryos were fixed and stained using 4’,6-diamidino-2-phenylindole, (DAPI) and fluorescent probes targeting *Ca*. F. symbiotica 16S rRNA and *Buchnera* 16S rRNA. Prior to *Buchnera* colonization, the syncytium of developing embryos is also devoid of *Ca.* F. symbiotica (Fig. 3A, SI1). The earliest observed entry into developing embryos was concurrent with *Buchnera* colonization (Fig. 3B, SI1). Older embryos contain *Ca*. F. symbiotica, where it is housed within bacteriocytes and sheath cells (Fig. 4C, 4D). Control images of uninfected aphids treated with the same probes and imaged with the same methods are included in SI2.

**FIG 4.**
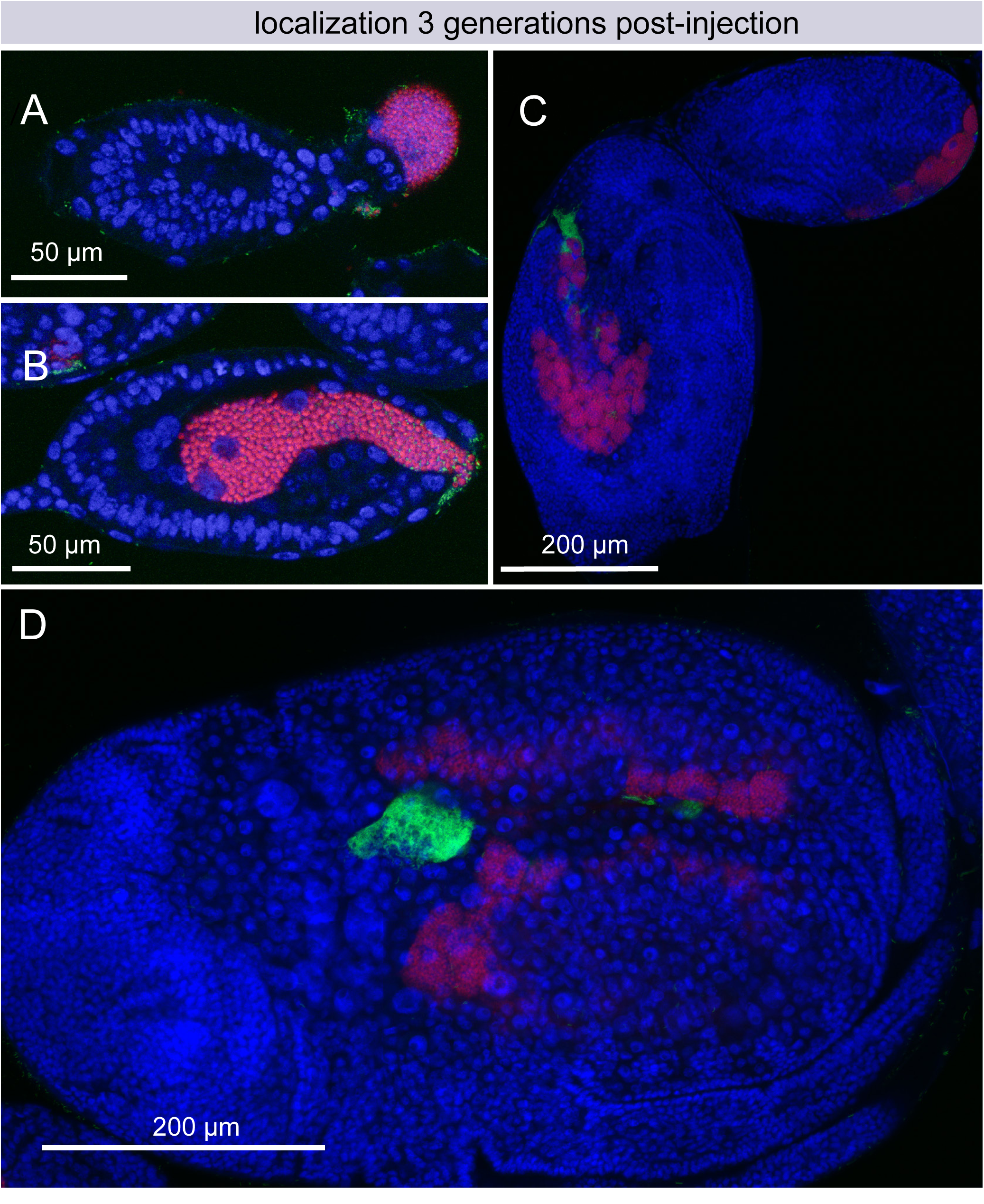
Localization of *Ca*. F. symbiotica in embryos three generations after injection. (A) Prior to *Buchnera* colonization, the syncytium of developing embryos does not contain *Ca. F. symbiotica*. (B) Entry of *Ca.* F. symbiotica into the syncytial space of stage 7 embryos is concurrent with *Buchnera* colonization. (C,D) *Ca.* F. symbiotica is visible in bacteriocytes and sheath cells of older embryos. *Ca.* F. symbiotica cells are labeled in green, *Buchnera* in red, and aphid DNA in blue. Linear adjustment for brightness was applied across all parts of each image to improve clarity. Two-channel images for FIG 4A, 4B are provided in FIG S2. Control images of aphid embryos not infected with *Ca*. F. symbiotica are shown in Fig. S1. Two channel images of FIG 4A and FIG 4B are shown in Fi. S2.

**FIG 5.**
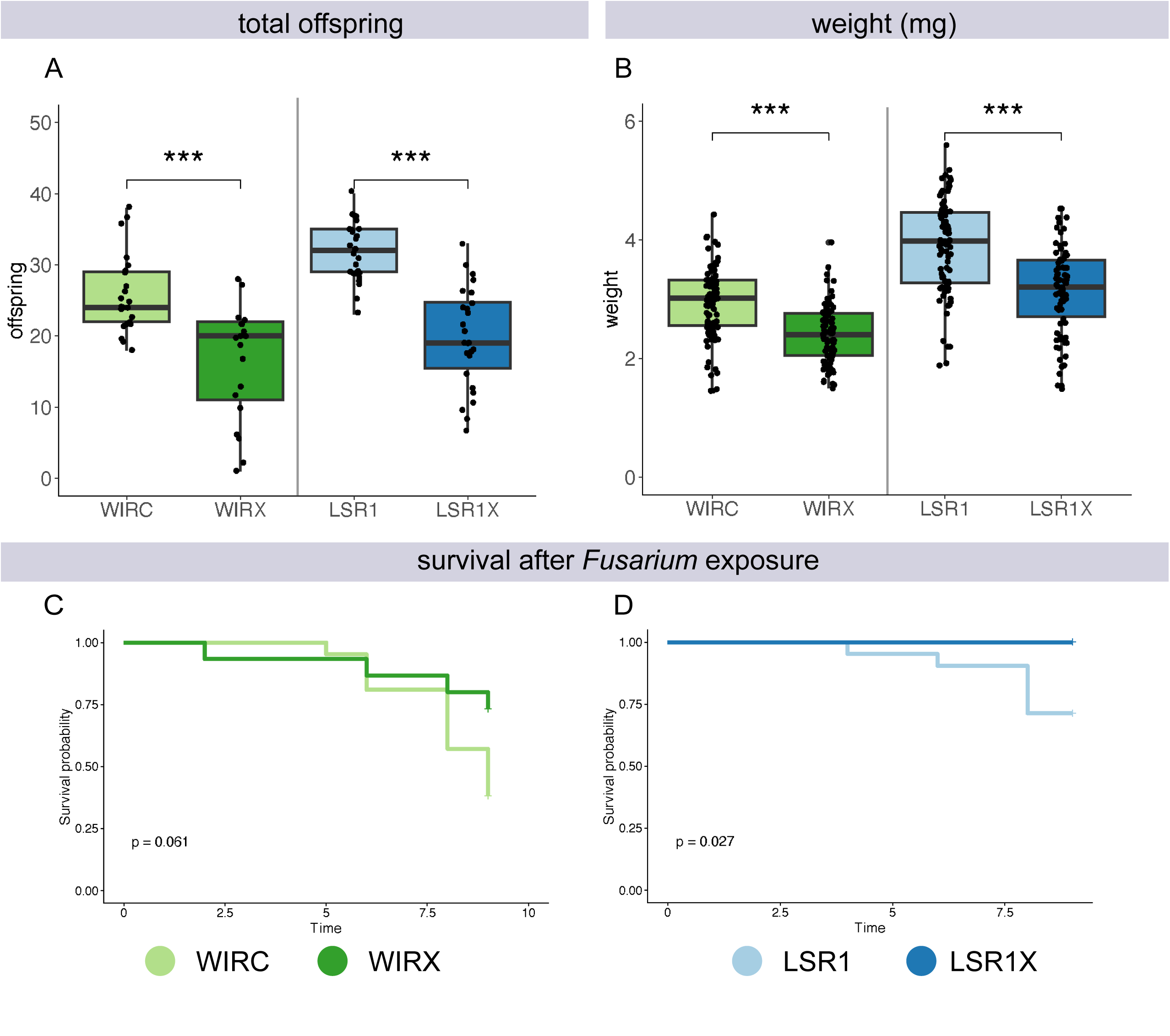
Phenotypic effects of *Ca.* F. symbiotica in native or artificial infections. (A) Boxplots indicating the number of offspring produced by aphids varying in infection status. ***, p<0.001, n = 22-27 aphids per treatment. (B) Adult mass measured in mg. ***, p<0.001, n = 82-96 aphids per treatment. Welch’s T-tests performed within each host genotypic background. (C-D) Kaplan-Meier survival curves of aphids after exposure to entomopathogenic *Fusarium.* (C) Mock treatments are depicted separately in S3 for clarity.

### *Ca*. F. symbiotica phenotypes are recapitulated in artificial infections, including protection against a fungal pathogen

High costs to host fitness have been associated with *Ca*. F. symbiotica infection under benign laboratory conditions (23, 25). To determine if our artificially infected lines exhibit similar effects, we measured the number of offspring produced within the first 5 days of reproduction and adult body weight. Artificially infected (LSR1X) and naturally infected (WIRX) lines were compared to uninfected aphids of the same host genetic backgrounds (LSR1 and WIRC, respectively). In both native or artificially generated infections, we observed a reduction in the total number of offspring produced. This reduction was statistically significant for both natural infections (t = 5.0382, df = 33.022, p < 0.001) and artificially generated infections (t = 7.6612, df = 40.164, p < 0.001). A significant reduction in adult body weight was also observed for aphids with native (t = 6.8436, df = 178.18, p < 0.001) and artificial infections (t = 6.3903, df = 165.86, p < 0.001).

To determine if *Ca*. F. symbiotica infection protects against additional non-specialist fungal pathogens, we performed pathogen challenges using a strain of *Fusarium* (Ascomycota), isolated from an aphid collected in Austin, TX USA. The same lineages of aphids described above were exposed to conidial suspensions or mock treatments of vehicle solution and mortality was monitored. Natively infected aphids experienced reduced mortality compared to uninfected aphids, although this difference was not statistically significant at α = 0.05 (Cox proportional hazard model, p = 0.061) (Fig. 4C). In the artificially infected line (LSR1X), this difference was significant (p = 0.027). Mock treatments experienced low mortality and did not differ by symbiont infection status (S3).

## DISCUSSION

*Ca.* Fukatsuia symbiotica, which is maternally transmitted but remains amenable to axenic culture, presents a rare opportunity to experimentally probe questions that are usually not feasible with vertically transmitted bacteria. Most vertically transmitted endosymbionts have lost functions required for living independently from their hosts, reflecting the gene loss that accompanies long term vertical transmission (30). One major advantage of independently cultivating symbiotic bacteria is the ability to efficiently sequence genomes without large portions of reads mapping to host DNA. Whole genome sequencing of *Ca*. F. symbiotica WIR places it among other *Ca*. Fukatsuia symbionts of aphids. *Ca*. F. symbiotica strains share average nucleotide identities >99.8%, even among distantly related *Cinara* and *Drepanosiphum* hosts. The high average nucleotide identity among these strains suggests relatively recent interspecific horizontal transmission, as previously noted (21). There is some evidence for intraspecific, horizontal transmission among pea aphids (18). Given the high sequence similarity shared among these strains, additional *Ca*. F. symbiotica strains from other aphid lineages may also be capable of independent cultivation.

Close relatives of the vertically transmitted aphid endosymbiont *Serratia symbiotica* also have been axenically cultured. However, they differ in fundamental features. These culturable strains of *S. symbiotica* comprise a clade of gut-associated pathogens that exhibit major differences in their gene content, localization, transmission, and effects on host fitness, and that differ from the vertically transmitted strains (28, 31–34). We found that *Ca*. F. symbiotica transmission occurs in a similar manner to observed for both culturable and non-culturable strains of *S. symbiotica* (28) with bacterial cells in the hemocoel entering embryos at an early stage of development when the syncytial cell that takes up *Buchnera* is exposed (Fig. 4B). Non-culturable *S. symbiotica* and culturable *Ca*. F. symbiotica WIR achieve stable long-term vertical transmission because they are less pathogenic enabling hosts to survive to become reproductive adults.

*Ca*. F. symbiotica strains have been shown to confer a variety of effects on host phenotypes (23). A strain collected in North America conferred protection against parasitoid wasps, when coinfecting with *Candidatus* Hamiltonella defensa (22). Strains of *Ca*. F. symbiotica from aphids collected in Europe are associated with many reported benefits, ranging from improved parasitoid resistance, defense against fungal pathogens, and improved recovery following heat stress (23, 24). However, at least one North American strain has failed to reproduce any of the same defensive phenotypes (25). Possibly methodological differences, including rearing temperatures, contribute to this variation in observed outcomes.

Culture-assisted methods may help clarify the impact of strain-level variation on the complex effects on hosts. Comparative genomics using protective and non-protective strains from pea aphids could provide insights into the genetic basis for these differences, and into potential mechanisms of protective effects.

We demonstrated that *Ca*. F.symbiotica strain WIR imposes substantial costs to host fitness in the absence of natural enemies, but also confers some protection against an entomopathogenic *Fusarium* isolate (Fig 4 C,D). Several other aphid endosymbionts including *Candidatus* Regiella insecticola, have been reported to protect against the distantly related aphid specialist *Pandora neoaphidis* (*35*). However, to our knowledge, no studies have shown pea aphid-associated heritable endosymbionts protecting against generalist entomopathogens, within Entomophthorales or Ascomycota (36, 37). Although *Ca*. Fukatsuia symbiotica imposes relatively high fitness costs under benign conditions, possibly the breadth of its protective effects helps explain its persistence in natural populations. In field collections, *Ca.* F. symbiotica commonly coinfects with *Ca.* H. defensa, and superinfection has been shown to reduce host fitness cost, which also helps explain *Ca.* F. symbiotica’s persistence in natural populations (25, 38).

Axenic culture may enable other approaches, including genetic manipulation. For example, the culturing of *Sodalis glossinidius* has made genetic methods possible for identifying genes involved in host colonization (39). Similar approaches may be applied in this system. Strain WIR grows slowly (∼14 days to observe colonies), as is consistent with intimate host association. Supplementation with additional B vitamins, amino acids, or sterilized aphid extracts failed to noticeably improve growth, and we saw no appreciable growth in the range of liquid media tested. Other groups have successfully used co-cultivation with insect cell lines for growing *Ca*. F. symbiotica, as well as *Ca*. Regiella insecticola and *Ca*. Hamiltonella in liquid media (21, 40, 41). Depending on the applications, a combination of these approaches may be useful for the study and manipulation of these endosymbionts outside of their hosts.

## MATERIALS AND METHODS

### Insect rearing

Pea aphid (*Acyrthosiphon pisum*) lines used for experiments were reared on fava seedlings (*Vicia fava*) at either at 15 °C for long term maintenance, or at 20 °C for microinjection experiments. Aphids were kept under long day conditions (18:6, L:D) to ensure clonal, asexual reproduction. The aphid line WIRX was collected in Madison Wisconsin, in 2017 and harbors a natural *Ca*. Fukatsuia symbiotica infection. This line was cured of a natural *Candidatus* Hamiltonella defensa infection in 2021. A subline was also cured of *Ca*. F. symbiotica infection as well, resulting in an aphid line cured of all secondary symbiont infection (WIRC). All experiments were performed more than 20 generations after antibiotic treatments. For recolonization and fitness experiments, the aphid line LSR1 was used and was previously cured of its natural *Candidatus* Regiella insecticola infection in 2009.

### Infection status

To screen for secondary symbiont infection, DNA was extracted from individual aphids using an ethanol precipitation protocol (42). Diagnostic PCRs were run using the primers 10F (AGTTTGATCATGGCTCAGATTG) and X420R (GCAACACTCTTTGCATTGCT) and performed using the cycling conditions described (17). 94 °C 2 min, 10 cycles of (94 °C 1 min, 65-55 °C in 1 °C steps every cycle 1 min, 72 °C 2 min) 25 cycles of (94 °C 1 min, 55 °C 1 min, 72 °C 2 min), 72 °C 6 min, hold 4 °C. PCR products were run on a 2% agarose gel, and visually assessed for amplification.

### Isolation of *Candidatus* Fukatsuia symbiotica

To grow *Ca.* F. symbiotica outside of its aphid host, single WIRX adults were briefly surface sterilized in 10% bleach with 0.5% TWEEN80, then rinsed in sterile water, before homogenization in 150 μL of phosphate buffered saline (PBS). This suspension was serially diluted in PBS and spot-plated onto HIA (Heart Infusion Agar) supplemented with 5% sheep’s blood, and Columbia Agar with 5% sheep’s blood. Plates were incubated under ambient atmosphere at room temperature (approximately 20 C) and wrapped in parafilm to prevent drying. Colony PCR was performed using the same primers for screening for *Ca*. F. symbiotica in aphids, and amplicons were submitted for Sanger sequencing at Eton Biosciences (San Diego CA, USA).

### Light and electron microscopy

Imaging was performed 14-21 days after inoculation onto fresh plates. For light microscopy, confluent *Ca*. F. symbiotica culture was suspended in PBS and imaged on a NIKON Eclipse TE2000-U epifluorescence microscope. For scanning electron microscopy, growth was fixed in 2.5% glutaraldehyde, and stained with 1.0% osmium in sodium cacodylate. Samples were incubated with 1.0% thiocarbohydrazide, before an additional treatment of 1.0% osmium in DI water. Graded ethanol washes were performed from 15% ethanol to absolute ethanol, and drying was performed after treatment in hexamethyldisilazane (HMDS). Dehydrated samples were sputter-coated in 5 nm gold/palladium and imaging was performed using a Zeiss Supra 40V Scanning Electron Microscope. Scanning electron microscopy (SEM) was performed at the Center for Biomedical Research Support Microscopy and Imaging Facility at UT Austin (RRID:SCR_021756).

### Whole genome sequencing and phylogenomics

Genomic DNA extraction and whole genome sequencing was performed by SeqCenter, Pittsburgh PA, USA. DNA extraction was performed using the Zymo Quick-DNA HMW MagBead Kit. Long reads were generated using Oxford Nanopore sequencing performed on R10.4.1 flowcells, run on a GridION platform. Short reads were generated using Illumina sequencing. Combined long and short reads were assembled using Unicycler (43). Small contigs under 200 bp were removed. Genome completeness and potential contamination was assessed using CheckM (44). Average nucleotide identity among *Ca*. Fukatsuia strains was calculated using FastANI (v1.34)(45). For phylogenetic placement select strains of interest were downloaded from NCBI (accessions in Table S2). Additional *Ca*. Fukatsuia genomes were accessed from Zenodo and were included in phylogenetic analyses (26). Assemblies were annotated with Prokka (v1.14.6)(46), and predicted proteomes were used as an input for OrthoFinder (v2.5.5) (47). A total of 340 orthologous proteins were aligned with MAFFT (v7.520) and concatenated using a custom script. Model selection was performed using ModelFinder (48) and a maximum likelihood phylogeny was generated using iqTree (v2.0.3), (49). Tree visualization was done using FigTree (v1.4.4).

### Host recolonization

To examine if *Ca.* F. symbiotica can recolonize its aphid host, confluent growth was scraped into PBS and diluted to an optical density at 600 nm (OD_600_) of 1.0. Approximately 0.1 uL of this bacterial suspension was injected into uninfected, 7 day old aphids of the LSR1 genotype. Injected aphids were transferred to fava leaves in petri dishes for 24 hr for recovery, before being transferred to plants. Once reproducing, aphids were transferred onto leaves in 1.5% agar daily. Offspring produced between 8-13 days after injection were sampled at 7-8 days old. Injections were repeated to assess stability of infections across generations. Here, groups of 7-10 injected aphids were transferred every two days. Mature offspring produced 10-12 days post-injection were placed on individual leaves in petri dishes in 1.5% agar and allowed to reproduce overnight. These aphids were screened for *Ca*. F. symbiotica infection as described above. Offspring from mothers testing positive were transferred to new plants. This process was repeated for another generation, and after the third generation post-injection, 6 sublines of infected aphids were maintained by transferring 3 adults to fresh plants. After 5 months, these lines were screened again for infection.

### Fluorescence microscopy of infected aphids

Fluorescence *in situ* hybridization (FISH) was performed using the protocol described in Koga et al (50). Offspring of aphids testing positive for *Ca.* F. symbiotica were dissected in 70% ethanol and embryos were fixed overnight in Carnoy’s solution (ethanol, acetic acid, chloroform 6:3:1). Fixed samples were dehydrated in a series of washes with 70% ethanol, followed by absolute ethanol. Samples were then bleached with 6% hydrogen peroxide in 80% ethanol for two weeks at room temperature, with the solution replaced every two days. Prior to hybridization, samples were washed with PBST (PBS, 0.2% TWEEN20). Hybridization was performed in hybridization buffer (20 mM Tris-HCl, 0.9 M NaCl, 0.01% sodium dodecyl sulfate, 30% formamide) containing DAPI and the probes ApisP2a-Cy5 (5′-Cy5-CCTCTTTGGGTAGATCC-3) targeting *Buchnera aphidicola* (*51*), and X16S-Alexa488 (5-Alex488N-CTCCATCAGGCAGATCC) designed in this study. Samples were mounted in SlowFade Gold Antifade Mountant (Invitrogen) and observed on a Zeiss 710 Confocal and Elyra S.1 structured illumination super resolution system. Probe specificity was confirmed by imaging uninfected aphid embryos (S2).

### Host fitness assays

Reproductive adults of each aphid line were placed on fava seedlings for 24 hours to generate age-controlled cohorts. Aphids were weighed on the first day of adulthood, transferred to individual fava seedlings, and monitored daily for the production of offspring. Reproductive output was measured as the number of offspring produced over the first 5 days of reproduction. Data for both reproductive output and weight are pooled from two trials initiated on separate days.

### Fungal isolation and pathogen challenge

To determine if *Ca*. F. symbiotica protects against ecologically relevant fungal pathogens beyond those previously reported, we attempted to isolate fungal pathogens from naturally infected aphids. Pea aphids were collected in Austin, Texas in April of 2023, and individual adults were placed on fava leaves in 1.5% agar. An individual aphid exhibiting fungal infection was ground in PBS and plated on Sabouraud agar (SDA). A single colony was passaged and maintained on yeast peptone, dextrose agar (YPD). For species identification, genomic DNA was extracted using a DNeasy DNA Extraction Kit (Qiagen), and a diagnostic PCR was performed, amplifying the internal transcribed spacer, using the primers ITS1 and ITS4 (52)n. Cleaned amplicons were submitted to ACGT (Germantown, MD) for Sanger sequencing. The consensus sequence for forward and reverse reactions was used as a query against the NCBI non-redundant nucleotide database to identify the fungal strain to genus.

For susceptibility assays, conidia were scraped from 14 day old plate cultures and suspended in 0.05% TWEEN80. Dosage was quantified using a hemocytometer and normalized to approximately 10^7^ conidia/mL. Aphids were each submerged in the conidial suspension, and transferred to fava seedlings. Mock treatments in which aphids were submerged in 0.05% TWEEN80 were also performed, and survival for each treatment was monitored daily.

### Statistical analyses

Statistical analyses were performed in RStudio, (R version 4.2.2). Comparisons of host fitness between infected and uninfected aphids were performed using Welch’s t-tests. Kaplan-Meier survival curves were generated using the survfit and ggsurvplot functions, in the R package survminer. Statistical significance was assigned at α = 0.05.

### Data availability

This Whole Genome Shotgun project has been deposited at DDBJ/ENA/GenBank under the accession JAXAWC000000000. The version described in this paper is version JAXAWC010000000. The associated BioProject Accession is PRJNA1043149.

## SUPPLEMENTAL MATERIAL

## ACKNOWLEDGMENTS

Light microscopy and SEM was performed at the Center for Biomedical Research Support Microscopy and Imaging Facility at UT Austin (RRID:SCR_021756). We thank Paul Oliphint, Michelle Mikesh, and Anna Webb for microscopy support. We thank the members of the Moran and Ochman labs at UT-Austin for discussions, input, and support. We thank Rachel Penczykowsk and Anthony Ives for the WIR aphid line. We thank Louis Deangelis for assistance in aphid collection.

We declare no competing interests.

This work was supported by a U.S. National Institutes of Health award R35GM131738 to NAM.

## Supplemental Figures and Tables

**FIG S1** Control images of aphid embryos not infected with *Ca*. F. symbiotica. Embryos were treated with the same hybridization solution as in Fig. 4, containing probes targeting *Ca.* F. symbiotica, *Buchnera*, and DAPI. Control embryos were imaged with the same exposure settings as in Fig. 4.

**FIG S2** Two channel images of FIG 4A and FIG 4B.

**FIG S3** Survival of mock treated aphids. Negative control aphids treated with 0.5% TWEEN80.

**Table S1** Genome assembly statistics for *Ca*. F. symbiotica WIR.

**Table S2** Genome accessions for strains used in Fig 2.

